# Arf GTPase-Activating proteins SMAP1 and AGFG2 regulate the size of Weibel-Palade bodies and exocytosis of von Willebrand factor

**DOI:** 10.1101/2021.03.29.437631

**Authors:** Asano Watanabe, Hikari Hataida, Naoya Inoue, Kosuke Kamon, Keigo Baba, Kuniaki Sasaki, Rika Kimura, Honoka Sasaki, Yuka Eura, Wei-Fen Ni, Yuji Shibasaki, Satoshi Waguri, Koichi Kokame, Yoko Shiba

**Affiliations:** Faculty of Science and Engineering, Iwate University, Morioka, Japan; Department of Molecular Pathogenesis, National Cerebral and Cardiovascular Center Osaka, Japan; Department of Biotechnology, National Kaohsiung Normal University, Taiwan; Department of Anatomy and Histology, Fukushima Medical University, Fukushima, Japan

**Keywords:** Arf, ArfGAP, secretory granule, exocytosis, vWF, WPB

## Abstract

Arf GTPase-Activating proteins (ArfGAPs) mediate the hydrolysis of GTP bound to ADP-ribosylation factors, which are important for intracellular transport. ArfGAPs have been shown to be critical for cargo sorting in the Golgi-to-ER and post-Golgi traffic. However, their roles in the sorting of secretory proteins remains unclear. Weibel-Palade bodies (WPBs) are cigar-shaped secretory granules in endothelial cells that contain von Willebrand factor (vWF) as their main cargo. WPBs are formed at the *trans*-Golgi Network, and this process is thought to be coupled with the sorting of vWF. WPB biogenesis was reported to be regulated by ADP-ribosylation factors and their regulators, but the role of ArfGAPs has been unknown. In this study, we performed siRNA screening of ArfGAPs to investigate the biogenesis of WPBs. We found two ArfGAPs, SMAP1 and AGFG2, to be involved in WPB size and vWF exocytosis, respectively. SMAP1 depletion resulted in small-sized WPBs, and the lysosomal inhibitor leupeptin recovered the size of WPBs. These results indicate that SMAP1 functions in preventing the degradation of cigar-shaped WPBs. However, AGFG2 downregulation resulted in the inhibition of vWF secretion upon Phorbol 12-myristate 13-acetate (PMA)-stimulation, suggesting that AGFG2 plays a role in vWF exocytosis. Our study revealed unexpected processes regulated by ArfGAPs for vWF transport.

**Summary Statement:** The ArfGAP proteins SMAP1 and AGFG2 were identified as regulating WPB size and vWF exocytosis.

## Introduction

Arf GTPase-Activating proteins (ArfGAPs) mediate hydrolysis of GTP bound to ADP-ribosylation factors (Arfs), small GTP binding proteins critical to the formation of transport vesicles(Kahn et al., 2008, Sztul et al., 2019, Tan and Gleeson, 2019). As ArfGAPs “inactivate” Arf-GTP by GTP hydrolysis, it had been thought that ArfGAPs were terminators of Arfs. However, there is an increasing amount of evidence to indicate that ArfGAP1, the first identified and most well-studied ArfGAP, plays an important role in cargo sorting during the formation of COPI vesicles(Pepperkok et al., 2000, Nickel et al., 1998, Lanoix et al., 2001, Yang et al., 2002, Liu et al., 2005, Nie et al., 2003, East and Kahn, 2011, Kahn, 2011, Spang et al., 2010, Beck et al., 2011, Hsu, 2011, Shiba et al., 2011). ArfGAP1 promotes the hydrolysis of Arf-GTP in the presence of coatomer, and its specific cargos, to promote the polymerization of coatomer (Luo et al., 2009, Shiba et al., 2011). ArfGAP1 could perform “shipping inspection” of cargos through its GAP activity during COPI vesicle formation (Shiba and Randazzo, 2012, Shiba and Randazzo, 2014).

The way in which ArfGAPs function in other sorting processes remains poorly understood. In endothelial cells, von Willebrand factor (vWF) is synthesized in the endoplasmic reticulum (ER), transported to the Golgi apparatus, then packaged into secretory granules called Weibel-Palade bodies (WPBs). Upon stimulation, WPBs fuse with the plasma membrane (PM) and vWF is released into blood vessels to recruit platelets for blood clotting. WPBs have characteristic cigar-shaped structures containing vWF as the main cargo. AP-1 is an Arf-dependent clathrin adaptor that mediates binding between clathrin and its cargo. Depletion of an AP-1 subunit results in the inhibition of biogenesis of WPBs from the *trans*-Golgi network (TGN) (Lui-Roberts et al., 2005). It was suggested that AP-1 and clathrin play roles in sorting vWF at the Golgi. However, other reports have suggested that AP-1 plays a role in the removal of lysosomal proteins from immature secretory granules (ISGs) in pancreatic β-cells, parotid acinar cells and neuroendocrine cells (Klumperman et al., 1998, Kakhlon et al., 2006), rather than the formation of ISGs. Therefore, the way in which AP-1 functions in both TGN and ISGs to sort each cargo remains unknown.

In this study, whether ArfGAPs regulate the sorting of vWF, we searched for ArfGAPs that regulate the formation of WPBs. First, we performed siRNA screening of ArfGAPs for WPB morphology using HEK293 cells, which have a high efficiency of transfection. Then, we used human umbilical vein endothelial cells (HUVECs) that have endogenous WPBs for a second screening. We identified two ArfGAPs, SMAP1 and AGFG2 which regulated WPB size and vWF exocytosis, respectively. Our study revealed the unexpected function of ArfGAPs in vWF transport.

## Results

### siRNA-screening of ArfGAPs in HEK293 and HUVECs

Our previous studies showed that depletion, but not overexpression, of ArfGAPs inhibits intracellular transport (Shiba et al., 2013, Shiba et al., 2010, Shiba et al., 2011). We therefore used the same approach to inhibit ArfGAP expression using siRNAs. There are 31 genes encoding ArfGAPs in humans. Of these, AGAP4 siRNA is able to target AGAP4 to AGAP10 mRNAs; we used a total of 25 siRNAs against ArfGAPs (Kahn et al., 2008, Shiba et al., 2013). For siRNA screening, we first used HEK293 cells, because HEK293 cells have a high efficiency of transfection, and the overexpression of the vWF gene in HEK293 is known to produce pseudo-WPBs (Romani de Wit et al., 2003, Michaux et al., 2003). We transfected the siRNAs of 25 ArfGAPs together with a GFP-vWF plasmid into the HEK293 cells and used confocal microscopy to examine whether the cigar-shaped structure of pseudo-WPBs was altered. In the first screening, we identified both the siRNAs of ArfGAPs that changed the cigar-shaped structure of pseudo-WPBs, and the siRNAs that we could not judge immediately as to whether the siRNAs affected the pseudo-WPBs structure. Fourteen of the 25 siRNAs against ArfGAPs remained. We again transfected the 14 siRNAs of ArfGAPs, and repeated the experiment. Six of the 14 ArfGAP siRNAs remained. The 6 ArfGAP siRNAs among 14 remained. To quantify the phenotype, we classified the cells based upon the structure of their GFP-vWF (Fig. 1A). Class I cells had cigar-shaped GFP-vWF, class II had small puncta of GFP-vWF, and class III had no apparent structure. We analyzed more than 30 cells per siRNA, and calculated the proportion of cells, which fell into each class. The cells transfected with the siRNAs of SMAP1, GIT2, AGFG2, ASAP2, ACAP3 and AGAP11 showed a decrease in class I cells, and increases in class II and III cells, compared with control siRNA-transfected cells (Fig. 1B).

**Figure 1.**
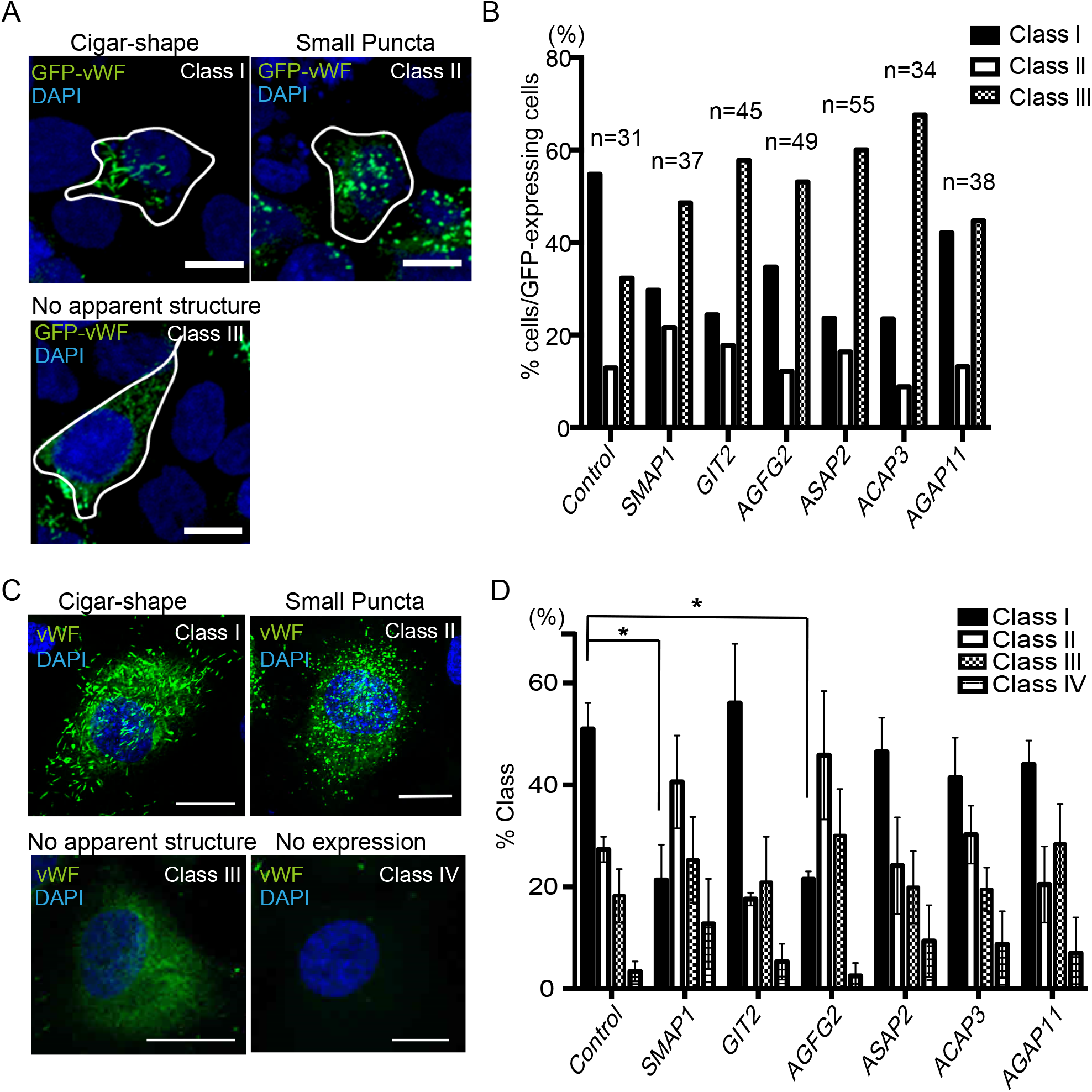
SMAP1 and AGFG2 were identified as affecting WPB morphology. **A**. HEK293 cells were transfected with GFP-vWF (green) and the nucleus stained with DAPI (blue). The images were captured using a confocal microscope, and the cells were classified into three classes. Scale bar, 10 µm. **B**. HEK293 cells transfected with the six siRNAs were classified into three classes and the percentage of each class in the GFP-expressing cells were calculated. More than 30 cells were analyzed, as described. **C**. HUVECs were stained with anti-vWF and DAPI. The cells were classified according to WPB structure. Scale bar, 15 µm. **D**. HUVECs transfected with the six siRNAs were classified into four classes, and the percentage of each class in all cells was calculated. More than 40 cells were classified, and the experiment was repeated three times (*n* = 3). One-way ANOVA followed by Dunnett’s multiple comparison tests were performed to compare each class with control cells. In SMAP1 and AGFG2 siRNAs-transfected cells, the number of cells in Class I was decreased. * *p* < 0.05, error bar; standard error of means (SEM).

Next, we tested whether SMAP1, GIT2, AGFG2, ASAP2 ACAP3 and AGAP11 siRNAs can affect the endogenous WPB structure in HUVECs. We electroporated these six siRNAs into HUVECs, stained them with anti-vWF antibody, and processed them for immunofluorescence. We classified the cells according to their WPB structure (Fig. 1C). Class I had cigar-shaped WPBs, class II had small puncta of WPBs, class III had no apparent structure, and class IV had no expression of vWF. We classified more than 40 cells in an experiment, and repeated the experiment three times. As shown in Figure 1D, the cells transfected with SMAP1 and AGFG2 siRNAs showed a decrease in class I cells compared with that in control cells (control 51%; SMAP1 21.4%; AGFG2 21.5%, *p* < 0.05 for both). These results suggest that SMAP1 and AGFG2 siRNAs affect the structure of endogenous WPB in HUVECs.

### SMAP1 depletion leads to small WPBs

To confirm the downregulation of SMAP1 and AGFG2 proteins, we performed immunoblotting using anti-SMAP1 and anti-AGFG2 antibodies for HUVECs electroporated with SMAP1 and AGFG2 siRNAs (Fig. 2A). We observed that SMAP1 and AGFG2 were downregulated by more than 70% and 80%, respectively. Hereafter, we refer to these SMAP1 and AGFG2 siRNA-transfected cells as SMAP1 knockdown (KD), and AGFG2KD cells, respectively. To quantify the changes in morphology of WPBs in SMAP1KD and AGFG2KD cells, we analyzed the size of the WPBs in immunofluorescence images (Fig. 2B and C). As shown in Fig. 2C, in SMAP1KD cells, the proportion of WPBs of length 0–0.5 µm and 0.5–1 µm were increased (0–0.5 µm: Control 45.4%; SMAP1KD 50.5%; *p* < 0.001; 0.5–1 µm: Control 22.5%; SMAP1KD 40.2%; *p* < 0.0001). WPBs of length 0.5–1 μm had a two-fold increase in the number of SMAP1KD cells. On the other hand, in SMAP1KD cells, the number of WPBs larger than 1 μm was decreased (1–1.5 µm: Control 12.5%; SMAP1KD 6.6%; *p* < 0.001; 1.5–2 µm: Control 6.8%; SMAP1KD 1.9%, *p* < 0.0001; >2 µm: Control 12.7%;

**Figure 2.**
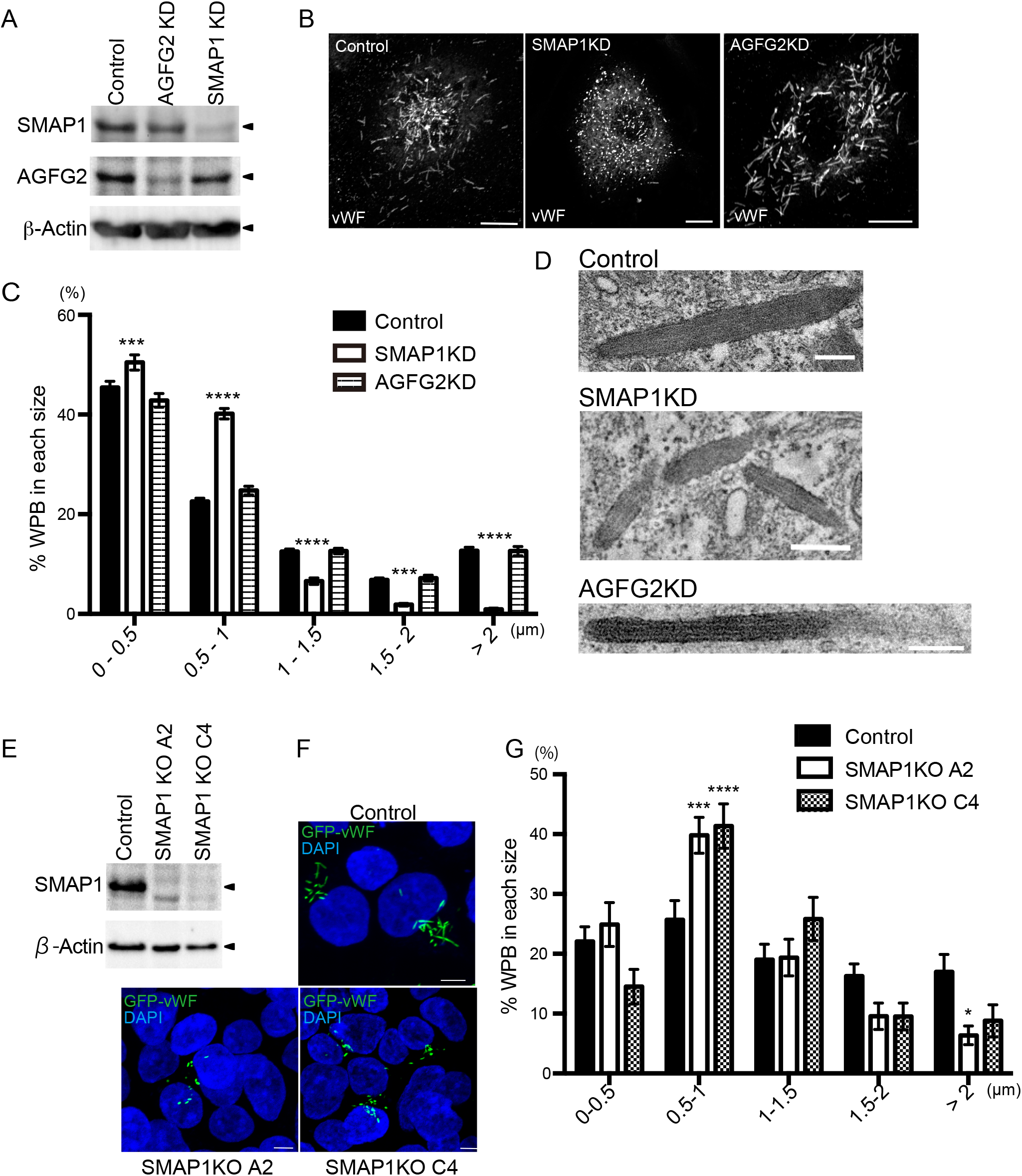
SMAP1 depletion results in small WPBs. **A**. HUVECs electroporated with control, SMAP1, or AGFG2 siRNA were subjected to immunoblotting with anti-SMAP1, AGFG2, and β-Actin antibodies. **B**. HUVECs electroporated with siRNAs of control, SMAP1, or AGFG2, were stained with anti-vWF. Scale bar, 10 µm. **C**. The size of WPBs of more than 10 cells was measured, and the experiment was repeated three times (*n* = 32 for all samples). The percentages of each WPB size are shown in the histogram. Two-way ANOVA followed by Sidak’s multiple comparisons test was performed for each siRNA. SMAP1KD cells have more WPBs less than 1 µm, and fewer WPBs larger than 1 µm. AGFG2KD cells do not show any difference from control cells. *** *p* < 0.001, **** *p* < 0.0001, ns; not significant, error bar, SEM. **D**. TEM images of control, SMAP1KD, and AGFG2KD cells are shown. In SMAP1KD cells, smaller WPBs were detected, while in AGFG2KD cells, cigar-shaped WPBs were observed. Scale bar, 200 nm. **E**. Control and SMAP1KO cell lines A2 and C4 derived from HEK293 were subjected to immunoblotting with anti-SMAP1 and anti-β-actin antibodies. **F**. Control and SMAP1KO A2 and C4 cells were transfected with GFP-vWF. Control cells have cigar-shaped pseudo-WPBs, while SMAP1 KO cells show small pseudo-WPBs. Scale bar, 5 µm. **G**. The experiment was repeated three times, and the size of pseudo-WPBs analyzed in more than 30 cells (Control, *n* = 42, A2; *n* = 39, C4; *n* = 40). Two-way ANOVA followed by Sidak’s multiple comparisons test was performed. SMAP1KO cell lines have smaller WPBs. * *p* < 0.05, *** *p* < 0.001,**** *p* < 0.0001, error bar, SEM.

SMAP1KD 0.9%; *p* < 0.0001). These results indicate that the WPBs were smaller in SMAP1KD cells. We performed the same analysis in AGFG2KD cells, and found no significant difference in WPB size (Fig. 2C). The reason why we could not reproduce the screening results in AGFG2KD cells is not clear. However, we noticed that the vWF signal in cigar-shaped WPBs in AGFG2KD cells was relatively weak. Anti-vWF antibody could have been difficult to be imported into WPBs in AGFG2KD cells (see Discussion). We used the same setting as in the control cells for quantification, so these parameters could have been too high for AGFG2KD cells. In that case, we could not have detected all cigar-shaped WPBs in AGFG2 KD cells during screening.

We further examined the structure of WPBs using Transmission Electron Microscopy (TEM). As shown in Fig. 2D, smaller WPBs were observed in SMAP1KD cells, whereas cigar-shaped WPBs were found in control and AGFG2KD cells.

To examine the off-target effects of SMAP1 siRNA, we established two cell lines of SMAP1 knockout (KO) by HEK293 cells. We did not use HUVECs, to avoid using the cells after passage 5. We confirmed complete depletion of SMAP1 protein by western blotting in A2 and C4 cell lines (Fig. 2E). We transfected GFP-vWF into the control, A2 and C4 cell lines, and analyzed the size of the WPBs (Fig. 2F and G). As in SMAP1KD cells, in A2 and 4 cell lines, WPBs sized between 0.5–1.0 μm were increased (control 24.5%; A2 39.2%; *p* < 0.001; C4 41.6%, *p* < 0.0001) WPBs >2 µm in length were decreased in the A2 cell line (control 16.5%; A2 5.6%, *p* < 0.05). The results confirmed that SMAP1 depletion causes smaller WPBs.

### vWF secretion was inhibited in AGFG2KD cells

To address the role of SMAP1 and AGFG2 in vWF secretion, we performed Enzyme-Linked Immunosorbent Assay (ELISA). We used Phorbol 13-myristate 12-acetate (PMA) as a secretagogue because it is highly efficient for stimulation (Zografou et al., 2012). We stimulated cells using 100 ng/ml PMA for 30 min, quantified the amount of vWF in the medium and the lysates, and calculated the percentage of secretion as the amount vWF in the medium divided by the total vWF in the medium and lysates. As shown in Fig. 3A, in control cells PMA stimulation increased vWF secretion around two-fold compared with PMA-unstimulated cells (control PMA-40.7%; PMA + 74.5%). Secretion without PMA is considered to be constitutive secretion. In SMAP1KD cells, the secretion of vWF with or without PMA showed no significant difference compared with that in the control cells. In contrast, PMA-stimulated secretion was halved in AGFG2 KD cells (control 74.5%; AGFG2KD 37.6%, *p* < 0.01). For PMA-unstimulated secretion, we observed some decrease in secretion in AGFG2KD cells, but the difference was not significant (control 40.7%; AGFG2 27.6%; *p* = 0.14). These results suggest that AGFG2 plays an important role in the PMA-stimulated secretion of vWF. We also quantified vWF secretion by immunofluorescence (Fig. 3B). In PMA-unstimulated cells, cigar-shaped WPBs were observed in the control cells. In contrast, PMA-stimulated cells have only small puncta, suggesting that most of the cigar-shaped WPBs were secreted upon stimulation. We quantified the size of the WPBs (Supplementary Fig. 1), and found that following PMA treatment the proportion of WPBs greater than 2μm in length was decreased (PMA-16%; PMA + 5.2%; *p* < 0.0001) and the proportion of WPBs between 0.5–1 µm was increased (PMA-30.7%; PMA + 38.6%; *p* < 0.0001). As the decrease of WPBs >2μm was marked following PMA treatment, with a decrease of 70%, we focused on quantifying WPBs>2μm. We performed immunofluorescence in AGFG2KD cells with or without PMA (Fig. 3B). In AGFG2KD cells, many of cigar-shaped WPBs remained, even following PMA treatment. We quantified WPBs >2μm (Fig. 3D), and found that AGFG2KD cells had around three-fold as many WPBs >2μm compared with control cells (control PMA + 5.9%; AGFG2KD PMA + 16.7%; *p* > 0.0001). Therefore, the immunofluorescence results were consistent with the ELISA results, supporting the idea that PMA-stimulated vWF secretion was inhibited in AGFG2KD cells. To check for off-target effects of AGFG2 siRNA, we overexpressed the siRNA-resistant AGFG2 gene with AGFG2 siRNA in HUVECs and performed immunofluorescence. As shown in Fig. 3C, AGFG2 overexpressed cells (AGFG2OE) lost cigar-shaped WPBs, even without PMA in control cells, and only cytoplasmic and perinuclear vWF remained. AGFG2 overexpression in AGFG2KD cells also led to the loss of cigar-shaped WPBs with or without PMA. We quantified WPBs>2μm and found that AGFG2OE cells had decreased WPBs>2μm in both control and AGFG2KD cells without PMA (Fig. 3D). PMA did not stimulate further vWF secretion. These results indicate that AGFG2 overexpression promotes vWF secretion and overcomes the AGFG2KD phenotype. Our analyses indicated that AGFG2 plays an important role in vWF secretion upon PMA stimulation.

**Figure 3.**
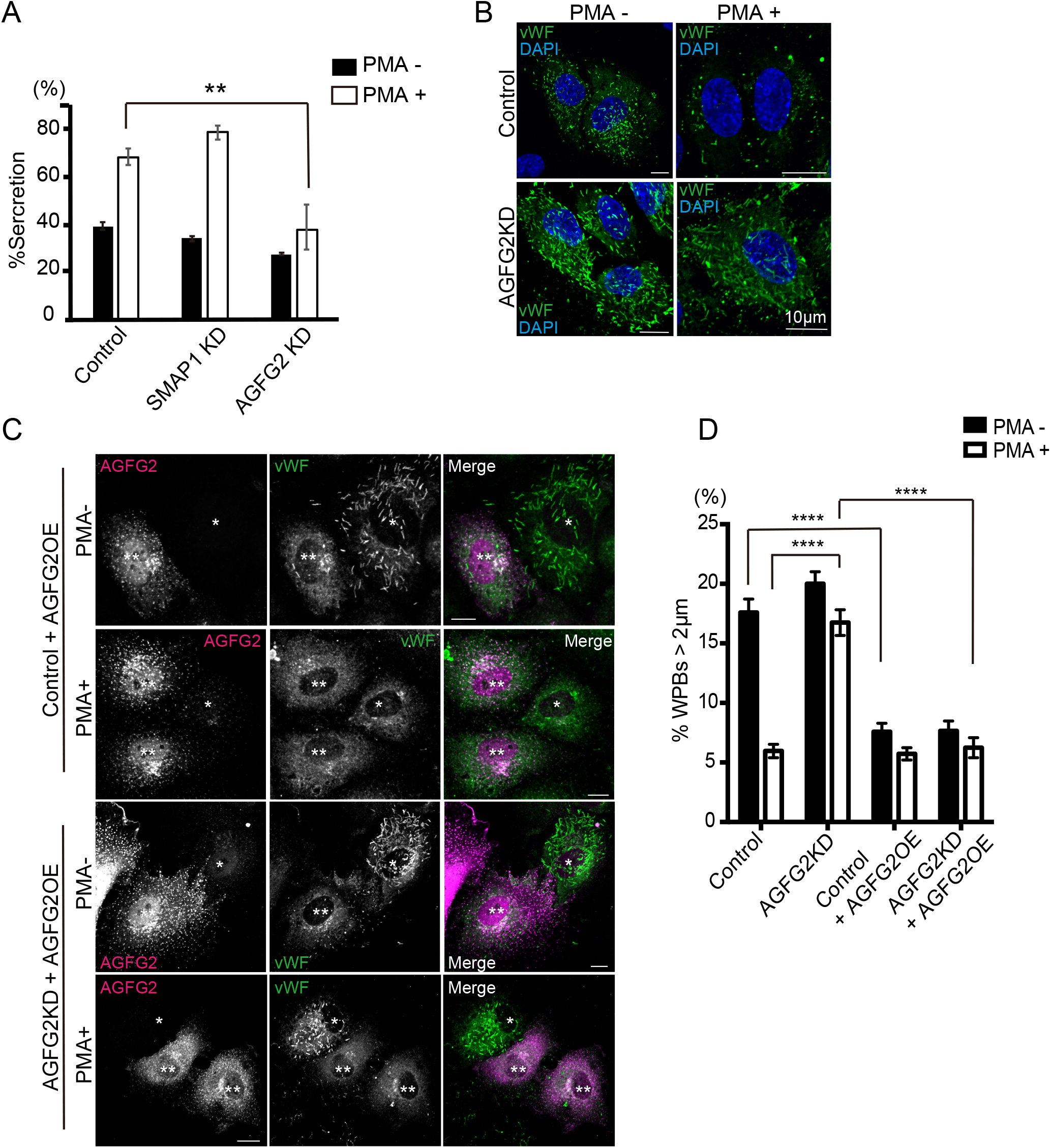
AGFG2 plays a role in vWF secretion. **A**. HUVECs electroporated with control, SMAP1, and AGFG2 siRNAs were stimulated with DMSO or 100 ng/ml PMA for 30 min, the medium and the lysates were collected, and the vWF amount was quantified using ELISA. The percentage vWF was calculated as the vWF in the medium divided by the total vWF. The experiments were repeated three times (*n* = 3). Two-way ANOVA with Dunnett’s multiple comparison test was performed with or without PMA. PMA-stimulated vWF secretion was significantly decreased in AGFG2KD cells. ** *p*< 0.01, ns; not significant, error bar, SEM. **B**. HUVECs electroporated with control and AGFG2 siRNAs were stimulated with DMSO or 100 ng/ml PMA for 30 min, and stained with anti-vWF (green) and DAPI (blue). Following PMA treatment, control cells lost cigar-shaped WPBs, whereas AGFG2KD cells retained cigar-shaped WPBs. Scale bar 10 µm **C**. HUVECs were electroporated with control or AGFG2 siRNAs, together with AGFG2 siRNA-resistant plasmids. The cells were treated with PMA for 30 min, and then stained for anti-vWF (green) and anti-AGFG2 (magenta) antibodies. AGFG2 overexpressing (OE) cells are indicated by **, and the cells without overexpression of AGFG2 are indicated by *. AGFG2OE cells lost cigar-shaped WPBs with or without PMA in both control and AGFG2KD cells. Scale bar 10 µm **D**. WPB size was analyzed for 15 cells per experiment, and the experiment was repeated three times (*n* = 45 for all samples). WPBs >2µm are shown. Two-way ANOVA with Sidak’s multiple comparison test was performed. **** *p* < 0.0001, error bar, SEM.

### WPB maturation was not highly perturbed in SMAP1KD and AGFG2KD cells

To investigate WPBs maturation in SMAP1KD and AGFG2KD cells, we examined the colocalization of vWF and the secretory granule marker, GFP-Rab27a. Rab27 has been reported to be a late stage marker for WPBs (Hannah et al., 2003). vWF was partially colocalized with GFP-Rab27a in control, SMAP1KD and AGFG2KD cells (Fig. 4A). We quantified the colocalization using Pearson’s correlation coefficient (PCC). The PCCs between vWF and GFP-Rab27a in control, SMAP1KD and AGFG2KD cells were 0.1 ± 0.08 (*n* = 14), 0.14 ± 0.23 (*n* = 10) and 0.26 ± 0.11 (*n* = 5), respectively. The values were all positive, confirming that GFP-Rab27a is recruited to WPBs in all cases. In SMAP1KD cells, WPBs were smaller but we did observe the colocalization of GFP-Rab27a with the small puncta of vWF (Fig. 4A). To examine whether vWF multimerization was affected, we stimulated HUVECs using PMA, and analyzed the multimerization of vWF in the medium (Fig. 4B). We found that in AGFG2KD cells, high-molecular weight (HMW)-vWF appeared to be decreased compared to that in control or SMAP1KD cells. We calculated the large-multimer ratio as the ratio of HMW-vWF to total secreted vWF, and compared each sample using the Large-Multimer Index (LMI, see Materials and Methods)(Fig. 4C). LMIs in control and SMAP1KD cells were not significantly different (control 311.6%; SMAP1KD 303.5%; *p* > 0.05), whereas AGFG2KD cells had lower HMW-vWF (control 311.6%; AGFG2KD 269.1% *p* < 0.01). These results suggest that vWF multimerization is inhibited in AGFG2KD cells but not in SMAP1KD cells. As the PMA-stimulated secretion of vWF was inhibited in AGFG2KD cells (Fig. 3A), lower levels of multimerized vWF could come from constitutive secretion. To further examine if WPB maturation was affected in AGFG2KD cells, we analyzed vWF processing in lysates by a protease furin that recycles between the TGN and the cell surface (Wagner et al., 1986, Wise et al., 1990, Molloy et al., 1994). The inhibition of the WPB maturation process would increase the pro-vWF amount. We observed mature vWF in SMAP1KD and AGFG2KD cells as well as in control cells (Fig. 4D). We quantified the percentage of mature vWF (Fig. 4E). Although we found that AGFG2KD cells and an approximately 9% decrease of mature vWF (control 83.2%; AGFG2KD 74.5%; *p* < 0.05), but most of the vWF matured in AGFG2KD cells. These results indicate that vWF maturation is not highly perturbed in SMAP1KD and AGFG2KD cells.

**Figure 4.**
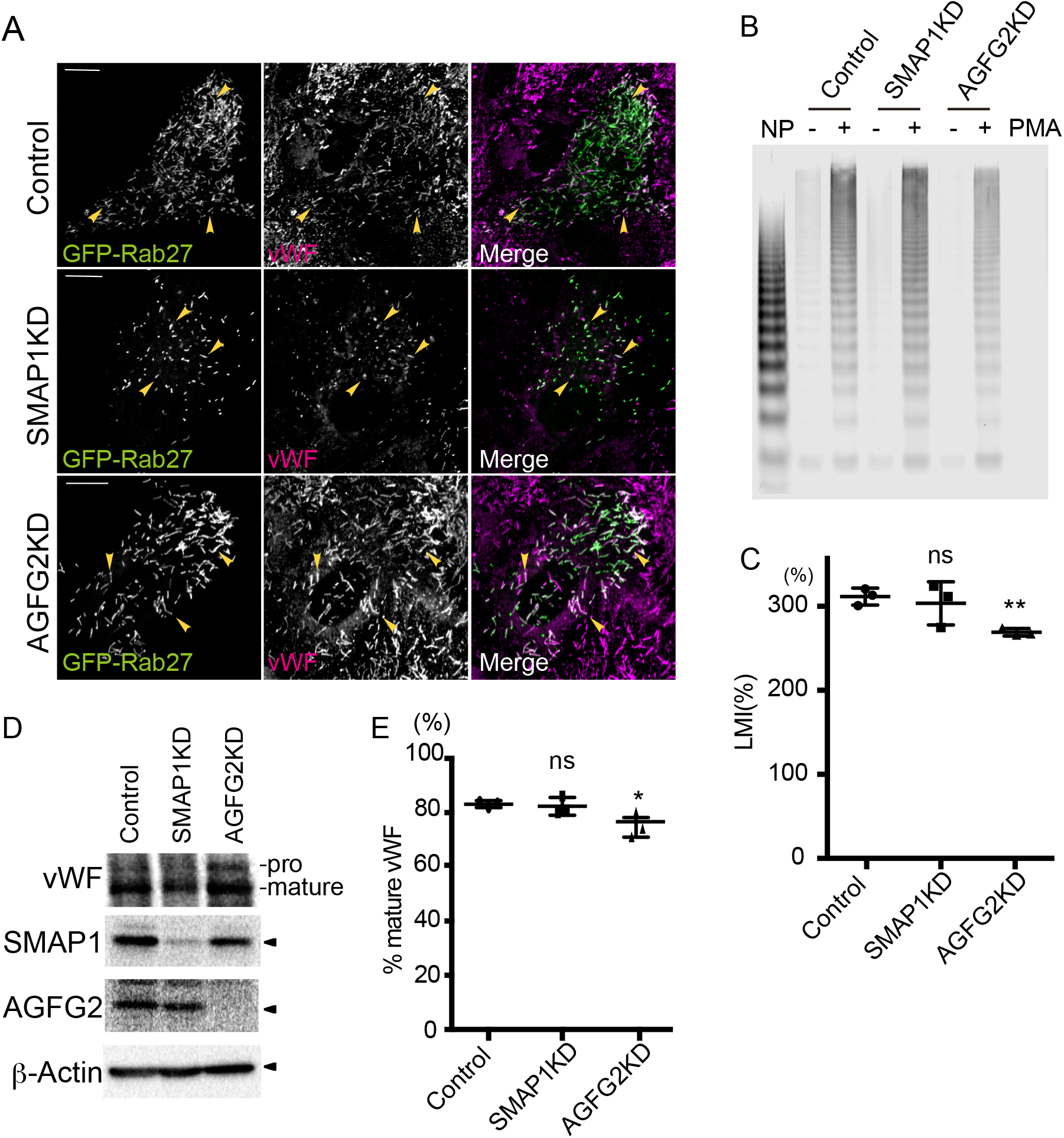
WPB maturation was not highly perturbed in SMAP1KD and AGFG2KD cells. **A**. HUVECs were electroporated with GFP-Rab27a and siRNAs, as indicated, and incubated for 72 hr. The cells were stained with anti-vWF (magenta). Arrowheads show colocalization between GFP-Rab27a and vWF. Small WPBs in SMAP1KD cells also colocalized with GFP-Rab27a. Scale bar 10 µm **B**. HUVECs electroporated with siRNAs as indicated were incubated for 72 hr and treated with 100 ng/ml PMA for 30 min. The medium was collected and subjected to agarose gel electrophoresis followed by anti-vWF western blotting. Normal human plasma (NP) was used as control. The smallest bands are considered to be dimers, and multiple bands are observed as multimerization increases. HMW-vWF was decreased in AGFG2KD cells. **C**. HMW-vWF was normalized by total multimers per lane, according to the large-multimer ratio. The LMI was calculated as the relative value of the large-multimer ratio to NP. The experiment was repeated three times, and unpaired *t*-tests was performed. AGFG2KD cells secreted less HMW-vWF, ns; not significant, ** *p* < 0.01, error bar, SD. **D**. HUVECs electroporated with siRNAs, as previously described, and the cell lysates were subjected to western blotting. Mature vWF was observed in all cells. **E**. The experiment in D was repeated three times, and the percentage of mature vWFs was calculated as the ratio to total pro- and mature-vWF. Unpaired *t*-tests were performed. A decrease of ∼9 % was observed in AGFG2KD cells, but most of the vWF matured, ns; not significant, * *p* < 0.05, error bar, SD.

To confirm that the vWF punctate signal does not come from other compartments, such as early endosomes, we stained vWF with the early endosome marker, EEA1 (Supplementary Fig. 2). vWF did not colocalize with EEA1 in control, SMAP1KD or AGFG2KD cells. The PCCs between vWF and EEA1 in control, SMAP1KD and AGFG2KD cells were −0.18 ±SD 0.07 (*n* = 5), −0.16 ± 0.07 (*n* = 5), and −0.02 ± 0.002 (*n* = 5). The values were all negative; that is, vWF was not mis-sorted to early endosomes. We also stained vWF with the TGN marker TGN46, and found that the TGN architecture was not significantly changed in SMAP1KD and AGFG2KD cells. These results indicate that SMAP1 plays a role in maintaining WPB size, and AGFG2 has a role in vWF secretion, independent of WPB maturation or vWF sorting at the TGN.

### Leupeptin recovered the size of WPBs in SMAP1-depleted cells

So far, our results indicated that AGFG2 plays a role in vWF secretion after the WPBs matured. However, it was not clear how SMAP1 regulates WPB size independent of WPB maturation or vWF sorting. We investigated whether SMAP1 regulates the degradation of cigar-shaped WPBs. We transfected GFP-vWF into the SMAP1KO cell line A2, and treated them with leupeptin, a soluble lysosome inhibitor that inhibits a wide range of proteases, including cysteine, serine and threonine proteases. We found that in leupeptin-treated cells, cigar-shaped WPBs re-emerged in SMAP1KO cells (Fig. 5A). We quantified the size of the pseudo-WPBs (Fig. 5B) and found that the numbers of WPBs >2µm was recovered by leupeptin (SMAP1KO 8.2%; SMAP1KO + Leup 17.8%; *p* < 0.01). To confirm these results for endogenous WPBs, we used HUVECs electroporated by either control or SMAP1 siRNA. We found that cigar-shaped WPBs were also restored with leupeptin treatment in HUVECs (Fig. 5C). Our quantification confirmed that WPBs >2µm was recovered by leupeptin (SMAP1KD 9.7%; SMAP1KD + Leup 13.9%; *p* < 0.01)(Fig. 5D). These results suggested that SMAP1 depletion accelerated the degradation of cigar-shaped WPBs, so only globular WPBs remained in SMAP1-depleted cells.

**Figure 5.**
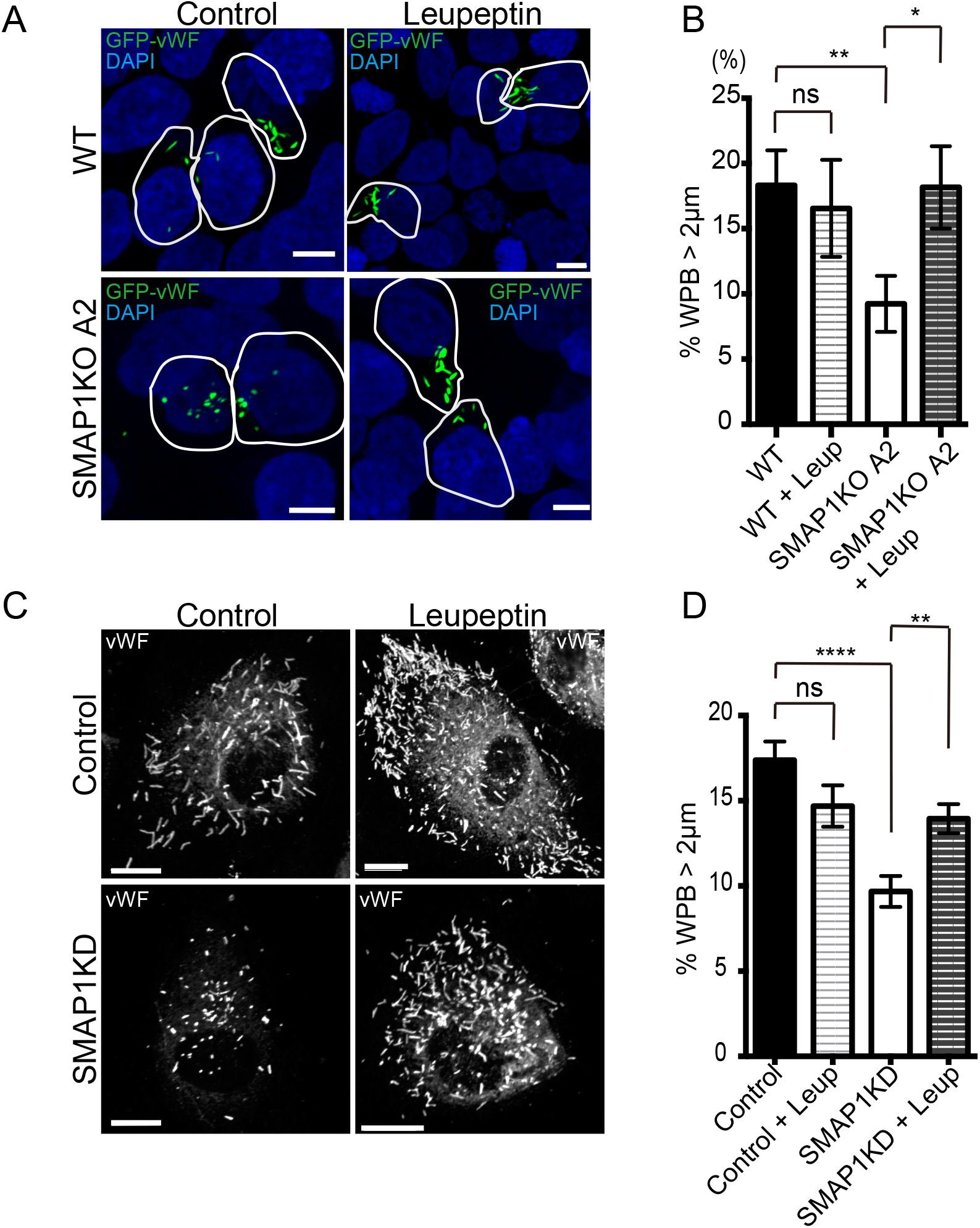
Leupeptin restored the size of WPBs in SMAP1-depleted cells. **A**. Wild-type HEK293 and SMAP1KO A2 cell lines were transfected with GFP-vWF, and 40 hr post-transfection, the cells were treated with 1 mg/ml Leupeptin for 8 hr. The cells were fixed and processed for immunofluorescence. SMAP1KO cells had small pseudo-WPBs, but leupeptin treatment recovered long pseudo-WPBs. Scale bar 5 µm. **B**. Pseudo-WPB size was analyzed for more than seven cells per experiment, and the experiment was repeated three times (WT; *n* = 35, WT + Leup; *n* = 25, SMAP1KO; *n* = 30, SMAP1KO + Leup; *n* = 39). Pseudo-WPBs >2µm are shown. Mann-Whitney tests were performed. The size of pseudo-WPBs was recovered by leupeptin in SMAP1KO cells. ns; not significant, * *p* < 0.05, ** *p* < 0.01, error bar, SEM. **C**. HUVECs were electroporated with control or SMAP1 siRNAs, and 60 hr post-transfection, 1 mg/ml leupeptin was added, and the cells incubated for 12 hr. The cells were fixed and processed for immunofluorescence. Cigar-shaped WPBs were recovered with leupeptin in SMAP1KD cells. Scale bar 10 µm. **D**. WPB size was analyzed for more than 10 cells per experiment and the experiment was repeated three times (Control *n* = 36, Control + Leup *n* = 33, SMAP1KD; *n* = 39, SMAP1KD + Leup; <u>n</u> = 34). WPBs >2µm was shown. Mann-Whitney test was performed. WPB size was recovered by Leupeptin treatment in SMAP1KD cells. ns; not significant, ** *p* < 0.01, **** *p* < 0.0001, error bar, SEM.

## Discussion

In this study, we identified SMAP1 and AGFG2 as ArfGAPs that regulate vWF transport. SMAP1 regulates the size of WPB, probably by inhibiting WPB degradation in the lysosome, and AGFG2 regulates the PMA-stimulated exocytosis of vWF.

Because AP-1 was proposed to be important for vWF sorting in the TGN(Lui-Roberts et al., 2005), we expected to find ArfGAPs involved in vWF sorting. We investigated the possibility that SMAP1 regulates WPB maturation, vWF multimerization, processing and localization. However, we could not find ArfGAPs involved in vWF sorting at the Golgi level in this study (Fig. 4 and Supplementary Fig. 2). Instead, we found that treatment with leupeptin restored the size of WPBs in SMAP1-depleted cells (Fig. 6). Therefore, we assumed that SMAP1 plays a role in preventing cigar-shaped WPBs from degradation in lysosomes, and depletion of SMAP1 leads to rapid degradation of cigar-shaped WPBs. WPBs are thought to be degraded by autophagy (Torisu et al., 2013, Wu et al., 2019), although vWF without a WPB membrane has also been seen in lysosomes (Torisu et al., 2013). Large sized organelles or pathogens could be preferably degraded by autophagy, but physiologically important organelles such as WPBs would escape from autophagy. The way in which SMAP1 prevents autophagy to protect WPBs from degradation, is an interesting research question.

To investigate whether the GAP activity of SMAP1 is important or not, we overexpressed wild-type SMAP1 and SMAP1[R61Q], a GAP-dead mutant of SMAP1 to HUVECs and HEK293 cells. The overexpression of SMAP1 and SMAP1[R61Q] both affected the Golgi architecture detected by TGN46, and vWF was accumulated in the ER (data not shown). Therefore, it was difficult to investigate the events after the TGN (Data not shown). To investigate whether SMAP1 localizes on WPBs, we examined the colocalization of endogenous SMAP1 and vWF outside of the Golgi area by immunofluorescence. We did not detect significant colocalization of SMAP1 on WPBs (PCC, −0.33 ± 0.12, *n* = 5). SMAP1 was very cytosolic in fixed cells. To determine the precise localization of SMAP1, we may need to develop the method by which SMAP1 is tagged in its genome and investigated in the endogenous level of the expression by live-cell imaging. Although Arf had been known to be very cytosolic, live-imaging of Arf using this method succeeded to detect Arf localization in transport intermediates (Bottanelli et al., 2017). The mechanisms how SMAP1 regulates WPB size, will be addressed in the future.

During the course of this study, Arf1, Arf4 and GBF1, a guanine-nucleotide exchange factor (GEF) for Arfs, were also reported to be important for vWF transport in the ER-Golgi level, but the phenotypes induced by the depletion of these factors are different to that of AP-1 depletion (Lopes-da-Silva et al., 2019). Considering these results, in conjunction with our own, we hypothesize that AP-1 is important to maintain the structure of the TGN, independent of the Arf-ArfGAP system.

The SMAP1 gene locus has been identified to be susceptible for pediatric venous thromboembolism (VTE)(Ruhle et al., 2017). That means physiologically, SMAP1 dysfunction promotes blood clotting. However, our results implicate that SMAP1 depletion causes small WPBs, which would lead to the secretion of less adhesive vWF to platelets, as small WPBs has been reported to secret vWF with lower adhesive activity to platelets (Michaux et al., 2006, Ferraro et al., 2016). Therefore, our results indicate that SMAP1 dysfunction would be predicted to cause bleeding disorder, but physiologically, the effects are opposite. Since SMAP1 is expressed in other cells including hematopoietic lineages (Kon et al., 2013, Kobayashi et al., 2014), VTE caused by SMAP1 dysfunction should be considered in a broader context, including the contribution of other cell types.

We also found AGFG2 plays an important role in vWF exocytosis. Initially, we identified AGFG2 as affecting the morphology of WPBs (Fig. 1). However, subsequent analyses showed that the size of WPBs was not changed in AGFG2KD cells (Fig. 2B, C and D). The reason that AGFG2 was identified by the screening of WPB morphology is not clear.

We noticed that, cigar-shaped WPBs in AGFG2 KD cells often have a weak immunofluorescence signal and could be difficult to detect, although they can be detected using higher gain settings in a confocal microscope. The weak staining of cigar-shaped WPBs in AGFG2KD cells could be due to the low accessibility of antibodies, as WPBs remain longer in AGFG2KD cells because of the lack of exocytosis, and vWF could be packaged more tightly in WPBs. As we used the same settings as those used to detect cigar-shaped WPBs in control cells for all images in the screening, a weak signal in AGFG2KD cells could go undetected, and only the small punctate structures of WPBs would be identifiable.

We detected the signal of AGFG2 in the Golgi area and the punctate signal probably near the cell surface by overexpression of AGFG2 (Fig. 3C). Because the overexpression of AGFG2 induced secretion of vWF, to investigate in which compartment AGFG2 colocalizes with vWF, we examined the colocalization between endogenous AGFG2 and vWF outside of the Golgi area. The PCC is −0.069 ± 0.18 (n = 10) without PMA and −0.14 ± 0.24 (n = 10) after PMA treatment for 30 min. we could not detect the significant colocalization between AGFG2 and vWF outside of the Golgi area. We also overexpressed AGFG2[R75Q], a GAP-dead mutant of AGFG2 to HUVECs. AGFG2[R75Q] expression also induced secretion of vWF, and we could not detect any difference on the effects compared with the expression of wild-type AGFG2 (data not shown). Human AGFG2 conserves ArfGAP consensus sequence CX2CX16CX2CX4R in ArfGAP domain, but AGFG proteins were reported to lose important amino acids in ArfGAP domain(Schlacht et al., 2013). Therefore, AGFG2 may not have the GAP activity originally in cells. AGFG2 also has FG repeat domain in its C-terminus. Previous studies showed that AGFG2 enhances the activity of HIV Rev for nuclear export of viral mRNAs (Doria et al., 1999). FG repeats are typical for nucleoporins that consist of nuclear pore complex, and known to play an important role in cargo selection in nuclear transport. Whether FG repeat or ArfGAP domain of AGFG2 is important for exocytosis, should be addressed in the future. According to the NCBI Gene database, AGFG2 is highly expressed in salivary gland (Fagerberg et al., 2014, Coordinators, 2016). AGFG2 could play a role in exocytosis in other secretory cells.

In this study, we identified SMAP1 and AGFG2 for regulating vWF transport. We revealed the novel roles of SMAP1 in regulating the size of WPBs and of AGFG2 in the vWF exocytosis. Future work will reveal the novel mechanisms of the regulation of organelle size by SMAP1 and exocytosis by AGFG2.

## Materials and Methods

### Reagents

ON-TARGET plus non-targeting siRNA #4, siGENOME SMART pool of human SMAP1, GIT2, AGFG2, ASAP2, ACAP3, AGAP11, and human AGFG2 gene were purchased from Horizon Discovery Ltd (Cambridge, UK). GFP-tagged human vWF plasmid (Romani de Wit et al., 2003) was kindly provided by Jan Voorberg (Stichting Sanquin Bloedvoorziening, Netherland) under Materials Transfer Agreement. AGFG2 was inserted into pcDNA3.1 using EcoRI and XbaI without any tag. For siRNA-resistant plasmids, AGFG2 was mutated in four locations with the primer, mutation #1; 5′-TAG TAT TTT TAC AAT CCC GTG GAA ATG AG-3’, 5′-ATT GTA AAA ATA CTA CTT CAG GCT CAG T-3′, #2; 5′-GGT TTG TAG AAA GAT TTG GTT GGG TCT G-3′, 5′-ATC TTT CTA CAA ACC TCA TTT CCA CGG GA-3′, #3; 5′-GAC AAG CCT CGT ACC AGA TTC CAG GGA T-3′, 5′-GGT ACG AGG CTT GTC CGA GCA TCA AAC AG-3′, #4; 5′-AGG AAG CGC GAA GTT GGG GCA GAG GCC A-3′, 5-AAC TTC GCG CTT CCT AAG TCC CCG AAG GA-3 using Prime STAR mutagenesis kit (Takara Bio Inc. Shiga, Japan). GFP-Rab27a was purchased from RikenBRC(Tsukuba, Japan)(Kuroda et al., 2002, Fukuda and Kuroda, 2002, Tsuboi and Fukuda, 2006). Sheep polyclonal anti-vWF antibody (AHP062T) was purchased from Bio-Rad (Hercules, CA, USA), anti-HA (HA.C5) from Abcam (Cambridge, UK), anti-EEA1(3C10) from MBL (Nagoya, Japan), anti-β-actin (8H10D10) from Cell Signaling (Danvers, MA, USA), anti-HA (12CA5) from Thermo Fisher Scientific (Waltham, MA, USA), rabbit polyclonal anti-SMAP1(A114714) and anti-AGFG2 (R08625) from ATLAS antibodies (Stockholm, Sweden), and anti-TGN46 (ab50595) from Abcam. Secondary antibodies of donkey anti-sheep, mouse and rabbit IgG conjugated with Alexa Fluor 488, 568 or 594 and Dylight were purchased from Thermo Fisher Scientific, AffinPure donkey anti-mouse and rabbit IgG conjugated with peroxidase from Jackson ImmunoResearch (West Grove, PA, USA).

### Cell culture

HEK293 cells were maintained by DMEM (Thermo Fisher) supplemented with 10% Fetal Bovine Serum (Cytiva, Marlborough, MA, USA) and Antibiotic-Antimycotic Stock Solution (NACALAI TESQUE, Inc., Kyoto, Japan). For replating cells, Trypsin-EDTA solution (Merck, Darmstadt, Germany) was used. HUVECs were purchased from Kurabo industries (Osaka, Japan) and Takara. HUVECs were maintained in Endothelial Cell Basal Medium 2 supplemented with Endothelial Cell Growth Medium kits (C22211, C22111, Takara). For replating cells, the cells were washed once with Hepes-buffered saline (Hepes-BSS), trypsinized with Trypsin-EDTA (CC5012, Lonza, Basal, Switzerland) and neutralized with Hepes-BSS/10% FBS. The cells were centrifuged to eliminate Hepes-BSS/10% FBS and plated with the growth medium. The cells were used by passage 5.

### siRNA screening of ArfGAPs in HEK293 cells

The custom cherry-pick siRNA library was constructed from 25 human ArfGAPs and control siRNA by Horizon Discovery Ltd. 15 × 10^4^ HEK 293 cells were seeded on collagen-coated coverslips (Cellmatrix, Type IV, Nitta gelatin, Osaka, Japan) and transfected with 5 nM siRNA using RNAiMax (Thermo Fisher) using a reverse-transfection protocol. The knockdown efficiency was confirmed by ArfGAP3 siRNA using western blotting, and almost 100% knockdown was achieved. After 24 hr of siRNA transfection, the medium was changed and the cells were transfected with GFP-vWF using Lipofectamine 3000, as per the manufacturer’s protocol (Thermo Fisher). The medium was changed after 24 hr and incubated for another 24 hr. The siRNA transfection period was 72 hr. The cells were fixed with 4% PFA/PBS for 15 min, quenched using 50 mM NH_4_Cl/PBS for more than 20 min, stained with DAPI (Merck) for 2 min, and mounted with Mowiol. A confocal microscope (Nikon C2, Tokyo, Japan) was used for capturing images using the 60x objective (NA1.40) and the 100x objective (NA 1.45). We visualized at all samples using the same setting of the confocal microscope, and eliminated any cells that had pseudo-WPBs with apparently no changes compared with those in control cells. We kept the cells for which we could not immediately judge whether there were any changes of pseudo-WPBs, and used them for the next analyses. Fourteen siRNAs (ArfGAP3, SMAP1, GIT2, AGFG2, ADAP1, 2, ASAP2, 3, ACAP1, 3, ARAP3, AGAP3, 4, 11) remained. We transfected these 14 siRNAs again into HEK293 cells, and analyzed them using the same protocol as the first screening. Six siRNAs (SMAP1, GIT2, AGFG2, ASAP2, ACAP3, AGAP11) remained. For these six siRNAs, the images of four fields that formed a single larger square were captured. More than 30 cells were analyzed (Fig. 1A and B).

### Immunofluorescence

Between 10–30 × 10^4^ HUVECs were electroporated with 50 pmol of control or ArfGAP siRNAs using the Neon transfection system (Thermo Fisher) at 1350V for 30 ms, seeded on gelatin-coated (Merck) coverslips. After 16–24 hr, the medium was changed. We noticed that the medium change promoted vWF secretion, therefore we incubated the cells for 48 hr after the medium change, to allow the cells to form new WPBs. Seventy-two hours post-transfection with siRNAs, the cells were washed in PBS, fixed in 4% PFA/PBS for 15 min, quenched using 50 mM NH_4_Cl/PBS for more than 20 min, permeabilized with 0.2% Triton X-100/PBS for 10 min and incubated with blocking solution (5% BSA/PBS) for more than 30 min. The cells were incubated with primary antibodies diluted in 0.02% Triton X-100/1% BSA/PBS for 90–120 min, washed with PBS for 5 min three times, and incubated with the secondary antibody diluted in 0.02% Triton X-100/1% BSA/PBS for 45 min. The cells were washed with PBS for 5 min three times, stained with DAPI for 3 min, washed, and mounted in Mowiol. The images were captured using a confocal microscope.

For Phorbol 12-myristate 13-acetate (PMA, Merck) treatment, HUVECs were electroporated and 48 hr after the medium was changed, the cells were serum starved for 1 hr, and then treated with 100 ng/ml PMA for 30 min, as described for ELISA. The cells were fixed and processed for immunofluorescence.

For leupeptin treatment, 15 × 10^4^ wildtype HEK293 cells or SMAP1 KO A2 cells were transfected with GFP-vWF with Lipofectamine LTX (Thermo Fisher) by reverse transfection, as per the manufacturer’s protocol. After 16–20 hr, the medium was changed. Twenty hours after medium change, leupeptin (Peptide Institute Inc. Osaka, Japan) was added to the medium to produce a final concentration of 1 mg/ml, and the cells were incubated for 8 hr. The cells were washed twice, fixed with 4% PFA/PBS, quenched with 50 mM NH_4_Cl/PBS, stained with DAPI and mounted with Mowiol. For HUVECs, 20 × 10^4^ HUVECs were electroporated with control or SMAP1 siRNA. For HUVECs, 20 x 10^4^ HUVECs were electroporated with control or SMAP1 siRNA. After 24 hr, the medium was changed. Thirty-six hours after the medium change, leupeptin was added to the medium to a final concentration of 1 mg/ml, and the cells were incubated for 12 hr. The cells were fixed and processed for immunofluorescence.

### Image analysis and statistics

Images of control, SMAP1KD, and AGFG2KD cells were captured using confocal microscopy. For quantification, the same settings were used in the control, SMAP1KD and AGFG2KD cells. To measure the size of WPBs, the confocal images were projected at maximum projection, and the WPB size was measured using the Fiji software (Schindelin et al., 2012). The “Analyze Particles” plugin was used to calculate Feret’s diameter as the length of WPBs. More than 10 cells per coverslip were analyzed and the experiment was repeated more than three times. More than 30 cells were quantified in total. A histogram was created using bins of 0.5 µm, as follows; 0–0.5 =< 0, 0.499>, 0.5–1 =< 0.5, 0.999>, 1–1.5 =< 1–1.499 >, 1.5–2 =< 1.5–1.999 >, 2< =< 2, infinity>. For PCC, a 3D mask was created using Fiji, and colocalization was quantified using the Coloc 2 plugin. The value was described with ±SD. Statistical analyses were performed using GraphPad PRISM version 6 software (GraphPad Software, San Diego, California USA, www.graphpad.com).

### Western blotting

180×10^4^ HUVECs were electroporated and seeded on a six-well plate. After 24 hr, the medium was changed and the cells incubated for 48 hr. The cells were lysed in 100 μl of lysis buffer (50 mM Tris-HCl, pH 7.5, 100 mM NaCl, 2 mM MgCl_2_, 1 mM DTT, 1% Triton X-100) with protease inhibitor cocktail (NACALAI) for 30 min on ice, centrifuged at 14,500 rpm for 10 min at 4°C, and the supernatants recovered. The protein concentration was measured using Bradford Protein Assay kits (Bio-Rad), and 30 μg of each protein was electrophoresed using 10% SDS-PAGE. The proteins were transferred to a PVDF membrane (Immobilon-P, Merck), blocked with 5% skim milk, and incubated with anti-SMAP1 and AGFG2 antibodies. The peroxidase-conjugated secondary antibody was incubated and reacted using ECL kits (Cytiva).The signal was detected using ChemiDoc XRS + (Bio-Rad). The membrane was stripped using stripping buffer (strong, NAKALAI) for detecting the β-actin signal as a loading control.

To investigate vWF processing, 50 × 10^4^ HUVECs were electroporated and seeded on a six-well plate. After 24 hr, the medium was changed and the cells incubated for 48 hr. The cells were scraped in 100 μl of lysis buffer, lysed by being drawn into and expelled from a 23 G needle five times, and kept on ice for 30 min. After centrifugation, the supernatant was taken and subjected to western blotting.

### Generation of SMAP1 knockout (KO) cell line

SMAP1 guide RNA was designed using the online design tool CRISPR direct (Naito et al., 2015). The selected gRNA targeted exon 2 of human SMAP1. The gRNA sequence was ordered as a pair of oligonucleotide 5′-CACCGATATTCCAGGAAGCCCATCG-3′, and 5′-AAACCGATGGGCTTCCTGGAATATC-3′. The oligonucleotides were annealed and inserted into the BbsI site of pSpCas9(BB)-2A-Puro(PX459)V2.0 (Addgene, Watertown, MA, USA), which has Cas9 and gRNA expression vectors. HEK293 cells were transfected with SMAP1-targeted PX459 vector using Lipofectamine 3000, incubated for two days, then 1 μg/ml Puromycin (Thermo Fisher) was added. After nine days, the cells were diluted to 1 cell /100μl medium, and 100 µl plated in each well of a 96-well plate. The cells were grown, and the genome was purified using QIAquick Gel Extraction kits (Qiagen, Hilden, Germany). A fragment of around 300 bp, including the SMAP1 PAM site, was amplified by the primer 5′-ACTGCCGCCAGTACCATTTG-3′, and 5′-TGGTAACGTCATGTTTACCTGTATC-3′, and sequenced by the nested primer 5′-TTGAGCATGTGACTTCTGTAAGC-3′. The SMAP1KO A2 cell line has a deletion of 14 bp after the PAM sequence, and the SMAP1KO C4 cell line has a T insertion in one allele, and an 11 bp deletion in a second allele. Both SMAP1KO cell lines were subjected to western blotting (Fig. 2E).

### ELISA

20 × 10^4^ HUVECs were electroporated with control, SMAP1 or AGFG2 siRNAs, and seeded on 24-wells plate coated with 0.3 mg/ml collagen type IV. After 16–20 hr, the medium was changed. Forty-eight hours after the medium change, the cells were washed twice with 0.1% BSA/ Hank’s Balanced Salt Solution (HBSS, Merck) and incubated for 1 hr with 0.1% BSA/HBSS. The cells were stimulated with either DMSO or 100 ng/ml PMA in 500 μl of 0.1% BSA/HBSS for 30 min. The medium was collected, centrifuged at 1000 rpm for 10 min at 4°C and stored at −30°C. The cells were washed with PBS twice, lysed with 500 μl of ELISA lysis buffer (0.5% Triton X-100, 1 mM EDTA/PBS) and stored at −30°C. Anti-vWF antibody was diluted in carbonate buffer (pH 9.6) at 1:400 for coating 96-well plates for 3 hr at 37°C. The coated plate was washed three times with washing buffer (0.1% Triton x-100/PBS), incubated with blocking buffer (0.1% Triton X-100, 0.2% gelatin /PBS) for 30 min at RT, and incubated with 180 μl of the sample and 20μl of blocking buffer (total 200 μl) overnight at 4°C. The plate was washed three times with washing buffer, and incubated with anti-vWF-HRP (P0226, Agilent technologies, Inc., Santa Clara, CA, USA) diluted 1:4000 in blocking buffer for 90 min at RT, then washed five times with blocking buffer. One tablet of 10 mg of O-Phenylenediamine dihydrochloride (Merck) was dissolved in 25 ml of citrate buffer, 200 μl added to each well, and the reaction started by adding 10 μl of 30% H_2_O_2_. After 3–4 min, the reaction was stopped with 50 μl of 2M H_2_SO_4_. The absorbance at 492 nm was measured using a plate reader (Powerscan HT, SD pharma, Osaka and Tokyo, Japan). The percentage of secretion was calculated by dividing the amount of vWF in the medium by the total amount of vWF in the medium and the lysate.

### TEM

For TEM, 25 × 10^4^ HUVECs were electroporated and plated on gelatin-coated coverslips. The medium was changed after 24 hr. Seventy-two hours post-transfection, the cells were fixed in 1% glutaraldehyde /4%PFA for more than seven days. The cells were quenched with 50 mM NH_4_Cl. The cells were treated with 1% osmium tetroxide, washed with PBS and 1% tannic acid for 30 min, dehydrated with ethanol, and embedded in Epok. Sections 100 nm thick were stained with U and Pb, and observed using electron microscopy (JEM2100, JEOL, Ltd, Tokyo, Japan).

### vWF multimer analysis

For multimer analysis, 15–20 × 10^4^ HUVECs were electroporated and plated on collagen-coated 24-well plates. After 16–20 hr, the medium was changed, and the cells incubated for another 48 hr. Seventy-two hours post-transfection, the cells were pretreated with 0.1% BSA/HBSS for 1 hr, then incubated with 0.1% BSA/HBSS including either DMSO or 100 ng/ml PMA for 30 min. The medium was collected, centrifuged at 1000 rpm for 10 min at 4°C and stored at −30°C. The experiments were performed in triplicate, and the media from all samples were electrophoresed on the same agarose gel. Normal human plasma (NP, Siemens Healthineers, Erlangen, Germany) was also electrophoresed, as a control. The gel was blotted on membrane and detected using anti-vWF primary antibody (Agilent DAKO, Carpinteria, California, USA) and IRDye 800CW-labeled anti-rabbit IgG secondary antibody (LI-COR Bioscience, Lincoln, NE, USA) as previously described (Naito et al., 2016). The near-infrared fluorescence signal was detected using an Odyssey CLx Imaging System (LI-COR Biosciences), and the intensity of each band was quantified using ImageJ (Schneider et al., 2012). The smallest five bands from smallest were considered to be “Low-Molecular Weight (LMW)-vWF,” bands 6 to 10 were “Middle Molecular Weight (MMW)-vWF,” and those more than 11 were “High-Molecular Weight (HMW)-vWF.” The intensity of HMW-vWF was divided by the total vWF intensity in the same lane to produce the large-multimer ratio. Then the LMI was calculated as the ratio of NP to the large-multimer ratio (Tamura et al., 2015).

## Acknowledgments

We greatly thank Dr. J. Voorberg for reading manuscript, Dr. Eiji Morita and Dr. Senri Ushida (Department of Biochemistry and Molecular Biology, Hirosaki University) for helpful discussions.

---

**Supplementary Figure 1.**
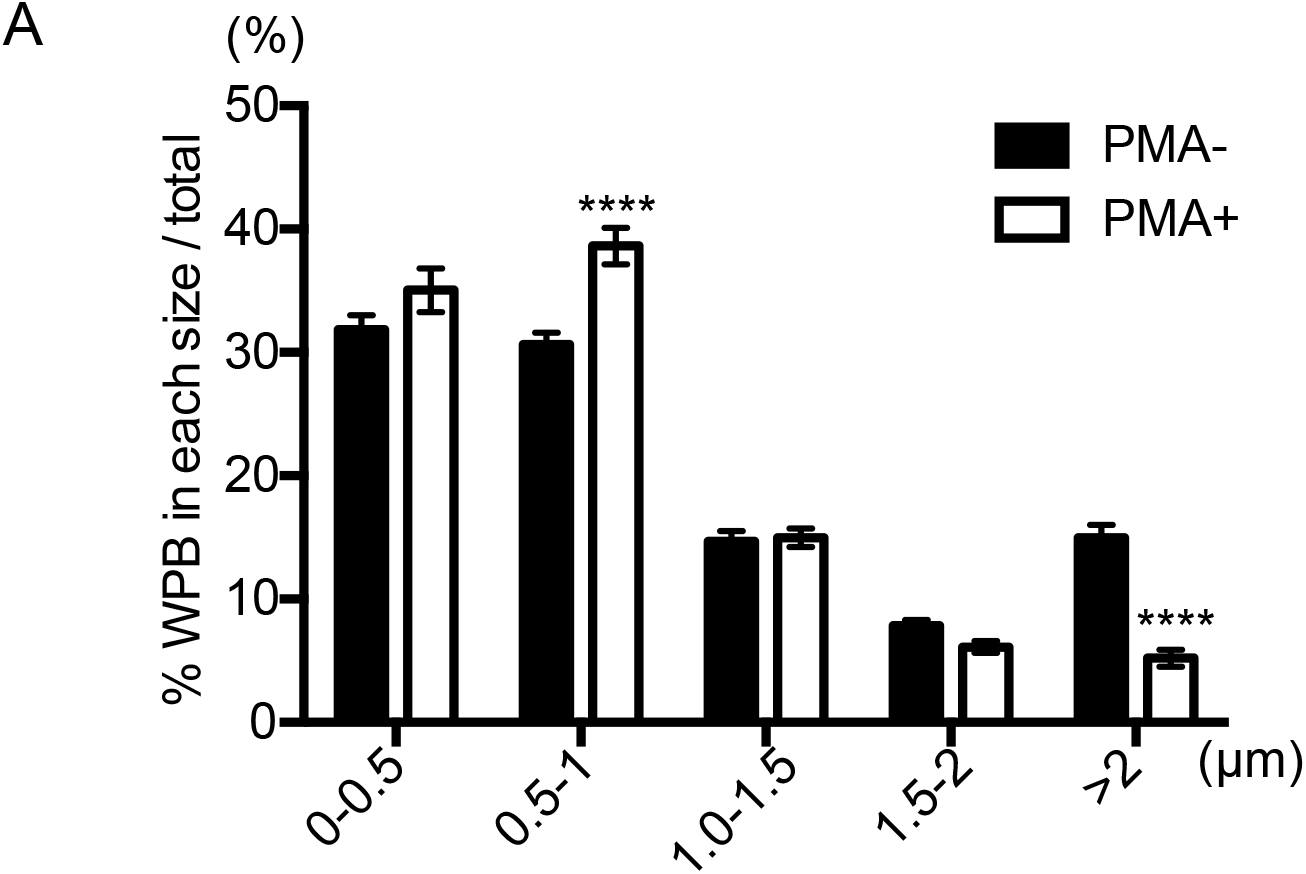
WPB size in HUVECs with or without PMA. HUVECs were plated in coverslips, and after 20 hr, the medium was changed. Forty-eight hours after the medium change, the cells were serum starved for 1 hr and treated by 100 ng/ml PMA for 30 min. The cells were fixed and stained for vWF. WPB size was quantified for 10 cells per experiment, and the experiment was repeated three times (*n* = 30 for both). Two-way ANOVA with Sidak’s multiple comparison tests were performed with or without PMA. WPBs >2 µm was markedly decreased by PMA treatment. **** *p* < 0.0001.

**Supplementary Figure 2.**
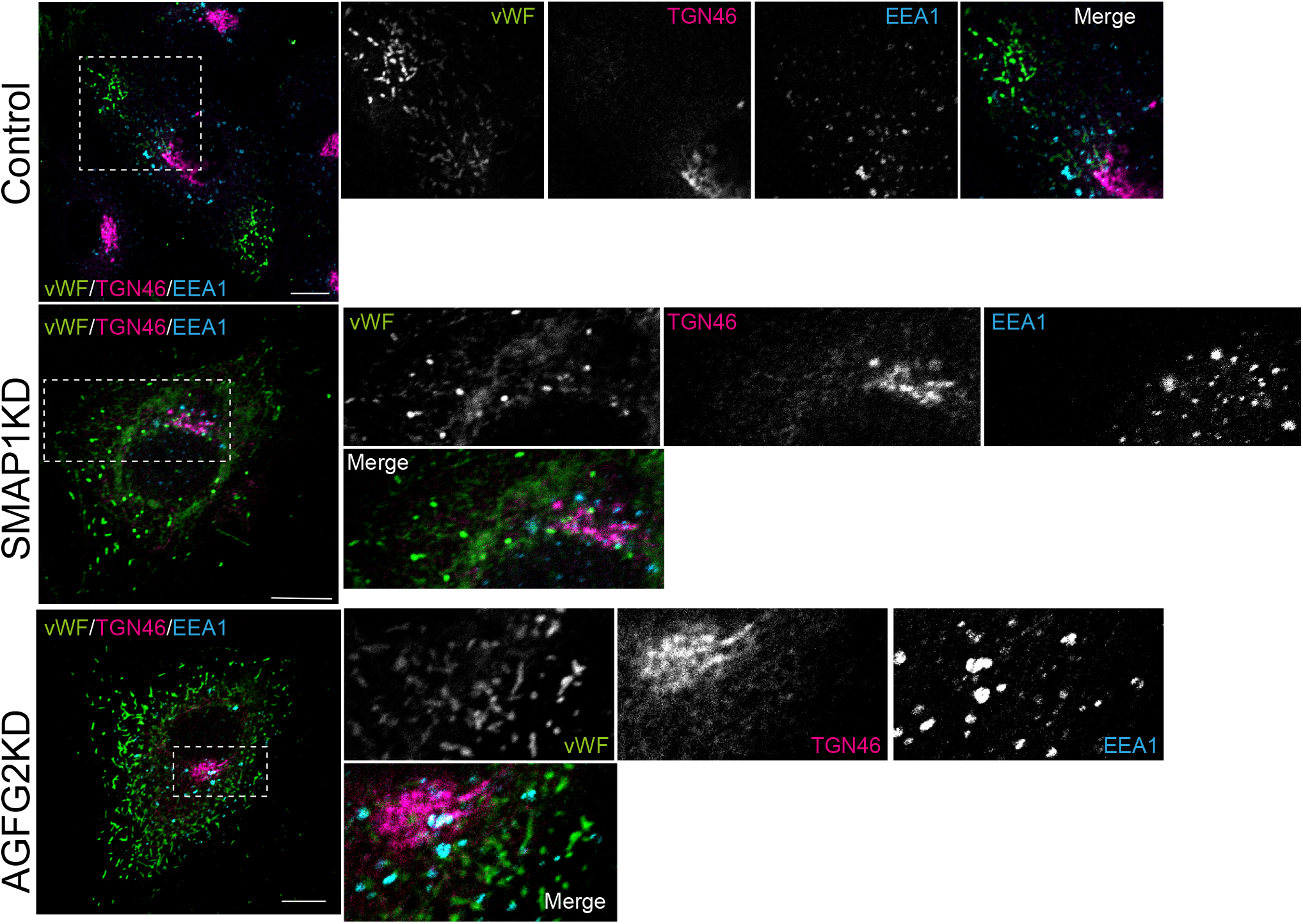
vWF did not colocalize with EEA1, and TGN architecture was not altered in SMAP1KD and AGFG2KD cells. HUVECs were electroporated with siRNAs as described, incubated for 72 hr, and stained for vWF (green), TGN46 (magenta) and EEA1 (Cyan). The area indicated by dashed boxes was enlarged. vWF did not colocalize with EEA1, and TGN architecture labeled by TGN46 was not altered in all cases. Scale bar 10 µm.

